# Attribution of Cancer Origins to Endogenous, Exogenous, and Actionable Mutational Processes

**DOI:** 10.1101/2020.10.24.352989

**Authors:** Vincent L. Cannataro, Jeffrey D. Mandell, Jeffrey P. Townsend

**Affiliations:** Department of Biology, Emmanuel College, Boston, MA; Program in Computational Biology and Bioinformatics, Yale University, New Haven, CT; Department of Biostatistics, Yale School of Public Health, New Haven, CT; Department of Ecology and Evolutionary Biology, Yale University, New Haven, CT

**Keywords:** cancer, tumor, single-nucleotide variants, mutational signatures, effect size, somatic mutation, selection, evolution, prevention, public health, molecular epidemiology

## Abstract

Mutational processes in tumors leave tell-tale genomic signatures composed of “passenger” mutations and mutations that have quantifiable effects on the proliferation and survival of cancer cell lineages. We identify the contributions of mutational processes to each oncogenic variant, quantifying responsibility for origination of changes at oncogenic variant sites contributing to tumorigenesis in 23 cancer types. We demonstrate that the variants driving melanomas and lung cancers are predominantly attributable to the actionable, preventable, exogenous mutational processes of ultraviolet light and tobacco exposure, whereas gliomas and prostate adenocarcinomas are largely attributable to endogenous processes associated with aging. Preventable mutations associated with pathogen exposure and APOBEC activity account for a large proportion of the cancer effect within head and neck, bladder, cervical, and breast cancers. These attributions complement epidemiological approaches—revealing the burden of cancer driven by single-nucleotide variants caused by either endogenous or exogenous, non-actionable or actionable processes, and crucially inform cancer prevention.

## Introduction

In the past half-century, our understanding of the origins of cancers has progressed from a deep mystery to a widespread acceptance that cancers are the outcome of an evolutionary process driven by mutation, consequent genetic variation, and selection on that genetic variation (Merlo et al., 2006; Nowell, 1976; Somarelli et al., 2020). Epidemiological studies have established both an association with age (Siegel et al., 2020) and causation by exposure to carcinogens (Smith et al., 2016), demonstrating that endogenous processes and exogenous mutagens can increase the rate of mutations (Barnes et al., 2018), create somatic genetic variation (Yates and Campbell, 2012), and increase the rate of cancer (Golemis et al., 2018; Greaves, 2015). In recent years, large-scale analyses of whole-exome and whole-genome tumor sequencing have been able to recover characteristic tissue-specific signatures of these underlying mutagenic processes in the patterns of variants that have suffused cancer genomes (Alexandrov et al., 2020). However, the specific cancer driver architecture within each kind of cancer tissue has also been demonstrated to be predictable (Hosseini et al., 2019) and, crucially, circumscribed (Venkatesan et al., 2017). Therefore, the causation of cancer by each mutational processes is not determined solely by their effect on mutation rate nor upon the amount of somatic genetic variation they induce, but critically depends upon the degree to which the specific mutations they supply provide selective advantages to clonal lineages within tissues that give rise to cancer.

To evaluate selective advantages requires knowledge of mutation bias, for which characteristic patterns have long been attributed to specific tissues (Brash et al., 1991; Pfeifer, 2015; Pfeifer et al., 2002; Poon et al., 2014). These patterns can be validated within model organisms under laboratory conditions and attributed to specific mutagenic sources (Segovia et al., 2015); numerous algorithms have been developed to deconvolve the total substitution load to its constituent mutational signatures (Grolleman et al., 2019). Application of these algorithms to whole-exome or whole-genome data recapitulates underlying mutation rates in their trinucleotide context without bias from natural selection because the vast majority of mutations are accumulating neutrally (Cannataro and Townsend, 2018; Greenman et al., 2007). Nevertheless, Poulos et al (2018) demonstrated that known major driver mutations are statistically associated with specific mutational signatures. Therefore, specific mutagenic processes in different tissues are driving tumorigenesis via mutations in genes that confer a survival and proliferative advantage to somatic cells. Estimation of the effects of each mutagenic process on the development of cancer requires quantification of the effects of each single nucleotide variant (SNV) toward tumorigenesis.

Quantification of the cancer effect of mutations requires estimation of their relative impact on cancer lineage survival and replication, an estimation that critically depends on an understanding of the baseline rate of mutation in the absence of natural selection (Cannataro and Townsend, 2018). Ostrow et al. (2014) performed a comprehensive analysis of ratios of non-synonymous change to synonymous change to quantify genomic natural selection in the somatic evolution of cancer. This approach has been applied in the field of evolutionary biology for decades and has recently been adapted to the nuances of cancer evolution in several meaningful ways (Shpak and Lu, 2016; Zhao et al., 2016), such as taking tissue-specific trinucleotide mutational patterns into account (c.f. Van den Eynden and Larsson, 2017). Martincorena et al. (2017) performed an analysis using trinucleotide substitution rates and covariate-informed gene-level mutation rates to quantify gene-wide selection conferring enhanced proliferation and survival of cancer cell lineages. Temko et al. (2018) deconvolved the underlying mutational signatures in tumor sets, associated signatures and drivers, and quantified the relative intragenic selection of the SNVs in a selection of high-burden driver genes. Cannataro et al. (2018) quantified the site-specific selective effect on each SNV during primary tumor development by determining the constituent mutational signatures driving mutation load in each tumor, coupling these rates with covariate-informed gene-level mutation rates, and quantifying their contribution to cancer cell lineage survival and reproduction in comparison to the convolved baseline mutation rate. These cancer drivers—and their relative effect—may be related back to the mechanisms driving genomic variation, i.e., the processes behind the detected mutational signatures.

Mutagenic environmental exposures have been correlated to specific cancer incidences by epidemiological studies spanning the previous 70 years (Doll and Hill, 1950; Loeb and Harris, 2008). Recently, cancer incidence has also been correlated with tissue-specific stem cell division numbers (Tomasetti et al., 2017; Tomasetti and Vogelstein, 2015), which has been interpreted as evidence that cancers are mainly driven by endogenous, i.e., aging or “bad luck”, effects. Other analyses dispute this conclusion, pointing out that it is confounded by the sensitivity of rapidly dividing tissues to exogenous mutational sources (Ashford et al., 2015; Wu et al., 2016), and by the exclusion of cancer types with known environmental causes (Wild et al., 2015). To determine the relative contributions of endogenous and exogenous processes on cancer phenotypes, tumor sequence data can be used to parameterize the magnitude of age-associated, exogenous and actionable mutational processes that contribute to molecular variation and the consequent cancer effects of each mutation attributable to these processes on tumorigenesis. Such analyses of the evolutionary dynamics driving tumorigenesis back to the sources of the heterogeneity fueling cancer evolution are essential to the advancement of our understanding of oncogenesis and cancer prevention.

Here we analyze the signatures of mutational processes in diverse cancer types. We quantify the cancer effect size of consequent single-nucleotide variants. We determine which cancer drivers in each tumor are attributable to actionable, and preventable, sources of mutagenesis. We quantify the contribution of each mutagenic process to cancer effect in individual patient tumors, and their relative contribution across tumors within sampled cancer types. We identify cancer types where the discrepancy between mutagenic input and cancer effect is largest, and smallest, and analyze which mutagenic processes are most proportionally discrepant with their cancer effect within each cancer type. This analysis enables comparison of the proportions of cancer effect attributable to age-associated processes to the proportions of cancer effect attributable to putatively preventable mutagenic processes such ultraviolet light exposure, tobacco smoking or chewing, and APOBEC mutagenesis, addressing a longstanding controversy regarding the role of endogenous “bad luck” and exogenous exposure to tumorigenesis—and moreover, informing the benefits of prevention of mutation in the prevention of cancer.

## Methods

To attribute the increased cellular reproduction and survival conferred by single nucleotide variants responsible for cancer growth to their underlying mutational sources we determined the sources of mutation within individual tumors, calculated the effect size of each single nucleotide substitution among tumors in each tumor type, and evaluated the likelihood that each of these substitutions was the product of each mutational source within each tumor. Thus, single-nucleotide substitutions responsible for the largest influence on cellular division and survival, and hence the cancer phenotype, may be attributed to the root sources of molecular variation within each somatic tissue. We analyzed the pan-cancer whole-exome tumor sequencing dataset curated in (Cannataro et al., 2018), except all Yale-Gilead tumors that might have been treated with chemotherapies were removed (removed tumors in **Table S1**). Scripts used to perform these analyses are available online (Townsend-Lab-Yale, n.d.).

### Attributing Sources of Mutation within Tumors

To attribute observed sets of substitutions in tumors to the underlying sources of mutations, we used the `R` package `deconstructSigs` (Rosenthal et al., 2016) to extract version 3.1 COSMIC mutational signatures from each tumor’s set of non-recurrent substitutions. We excluded recurrent variants because they are much more likely to be under selection in the cancer cell population; non-recurrent mutations more accurately reflect mutational influx. To minimize signature bleeding because some COSMIC signatures share similar mutational profiles, we limited the number of signatures detectable in each tumor type to those signatures detected at any prevalence in tumors of that type previously by Alexandrov *et al.* (2020), with the addition of enabling inference of SBS16 within esophageal squamous cell carcinoma (Li et al., 2018). We also applied the recommended minimum threshold for the number of substitutions necessary to attribute to a signature associated with increased mutagenesis. For example, signatures attributable to defective DNA mismatch repair were only allowed in tumors with over 200 substitutions (Alexandrov et al., 2020). Some tumors analyzed exhibited fewer than 50 substitutions (**Supplemental Fig. S1**)—a threshold below which precise deconvolution of mutational signatures becomes problematic (Rosenthal et al., 2016). For these tumors, we mixed the `deconstructSigs` estimates of the signature weights for the specific tumor with the average signature weights for the tumors with 50 or more substitutions of the same tumor type, weighting the former in proportion to the number of variants in the tumor out of 50.

As some COSMIC signatures have been attributed to artifactual processes such as sample handling and sequencing, we focus on the tumor-type-specific subset of signatures *B* that represent biologically relevant mutational processes (Alexandrov et al., 2020). The fitted weights of signatures in *B* reflect the relative rates that their underlying mutational processes contribute mutations. To determine the tumor-specific relative weight *w_i_* of a biological signature *i* ∈ *B*, we divided its fitted weight 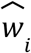 by the sum of the fitted weights of all biologically associated signatures; i.e.,

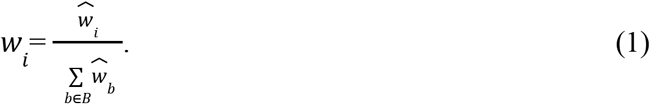

### Calculating the mutational source variant weight

After calculating the tumor-specific relative weights of mutational sources (**Fig. 1A**) as described in **Eq. 1** (**Fig 1B**), we used the trinucleotide-context-specific relative mutation rates defined for mutational signatures to calculate the probabilities that each variant in each tumor derives from the mutational processes underlying each biological mutational signature. Let *ψ* be a matrix of trinucleotide-context-specific relative mutation rates for the signatures in *B*, with *ψ_i,j_* being the rate for signature *i* of trinucleotide-context-specific mutation *j*. Retrospectively, the probability that a single-nucleotide variant constituting a trinucleotide-context-specific mutation *j* derived from the process underlying mutational signature *i* in tumor *n* is

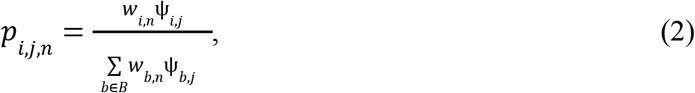

where each *w_k,n_* is the relative weight of signature *k* in tumor *n* (**Eq. 1**).

**Figure 1:**
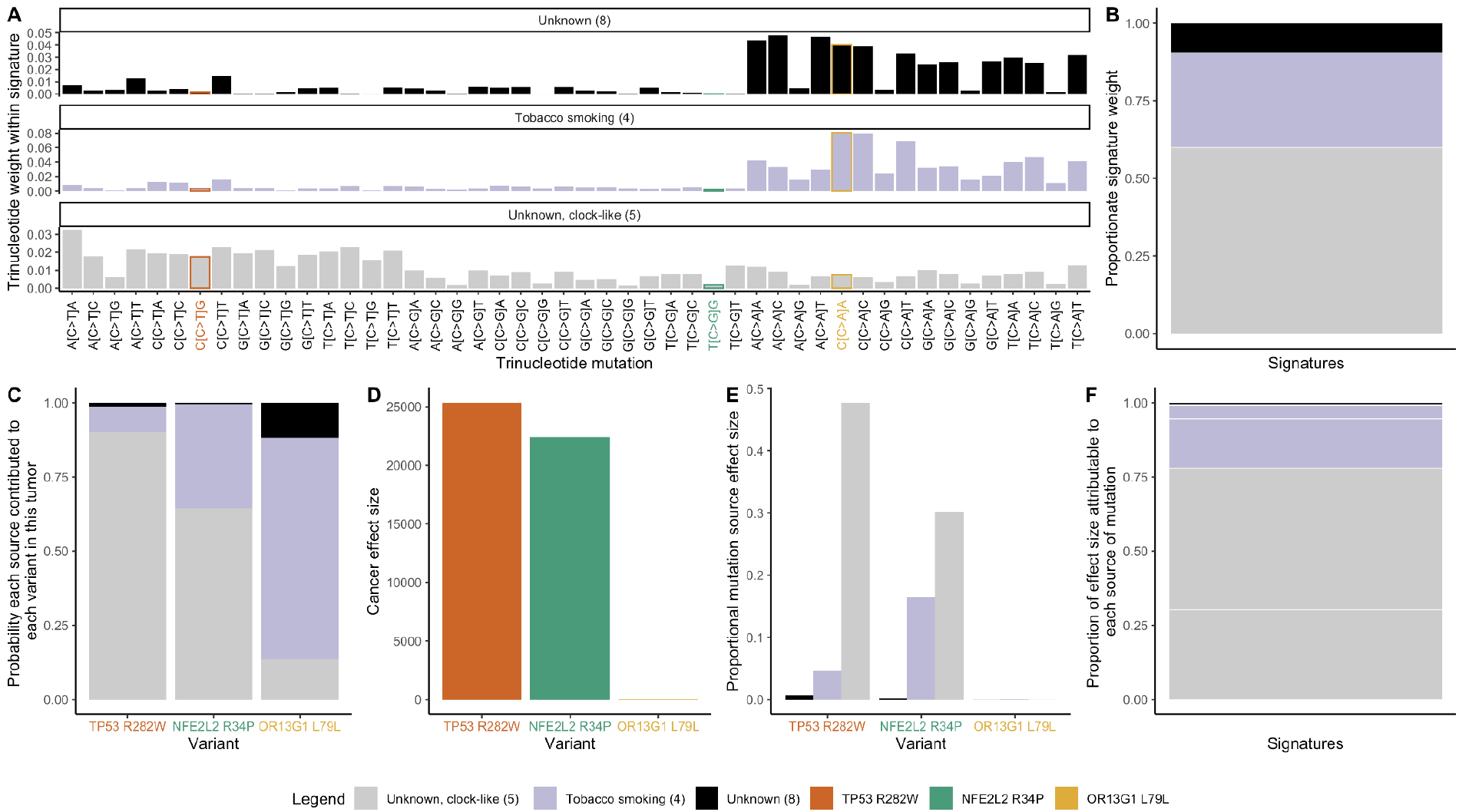
Calculating the cancer effects by mutational source. The respective products of (**A**) the trinucleotide weights within each signature (Alexandrov et al., 2020), e.g. a relatively infrequent CCG→CTG mutation leads to TP53 R282W, orange; a relatively infrequent TCG→TGG mutation leads to NFE2l2 R34P, green; and a relatively frequent CCA→CAA mutation leads to OR13G1 L79L, tan; and (**B**) the proportions of observed mutations in a tumor caused by each signature (from deconstructSigs, Rosenthal et al., 2016) (gray: unknown but age-related; lavender: tobacco smoking; black: unknown signature 8) can be normalized to yield (**C**) the probability each source contributed to each variant in this tumor. Each of these probabilities serves as a weight to multiply by (**D**) the cancer effect size of each variant (Cannataro et al., 2018), to yield (**E**) the probability-weighted portion of effect size for each variant attributable to each source of mutations, stacked to compose (**F**) the proportion of cancer causation attributable to each source of mutations. This example illustrates the calculation on a lung squamous-cell carcinoma, MDA-1229-T.

For a given variant, we can scale these probabilities by the impact of the variant on the survival and proliferation of cancer cells—that is, the variant’s cancer effect size—to quantify the relative contributions of each mutational source to the cancer.

### Calculating the Effect Sizes of Variant Substitutions in Tumors

To attribute a cancer effect to each substitution, we used the R package cancereffectsizeR, version 2.1.2 (Cannataro et al., 2018). As described in our previous work, the package’s underlying model assumes that substitutions fix in accord with a Poisson distribution at the rate that mutations arise (mutation rate μ) multiplied by their cancer effect size γ. The term γ is 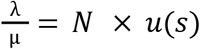, where λ is the rate of substitutions, *N* is the effective population size of cancer cells, and *u*(*s*) is the probability of fixation of a new mutation as a function of the selection coefficient *s*, from population genetic theory. Thus, for every variant, we calculated the effect size that maximized the likelihood function

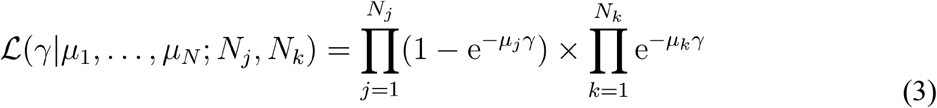

for the *N* tumors, *N_j_* of which exhibited at least one substitution event of that variant in tumor *j* ≤ *N*, and *N_k_* of which exhibited no such variant in tumor *k* ≤ *N*. Each tumor-specific mutation rate *μ*_1_, …, *μ_N_* was calculated by extracting the mutation rate in each trinucleotide context of each variant from the tumor-specific mutational signature weights (**Eq. 1**) and convolving it with the gene-specific mutation rate as in Cannataro et al. (2018).

### Calculating mutational source effect weight

The relative contribution to cancer effect of variant *i* from mutational process *b* in tumor *n* ≤ *N*,

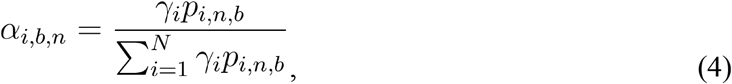

scales the effect size *γ_i_* by the probability it was caused by the process underlying signature *b*, within each tumor, relative to all contributions of cancer effect in that tumor.

To quantify the contribution of each mutational process to the total relative cancer effect of a variant in tumors of each cancer type, we average *α_i,b,n_* across all *N* tumors for a fixed value of indices *b* and *i*. To quantify the proportion of population-level burden of cancer effect size contributed by each mutational process to each tumor type, we sum *α_i,b,n_* across all *i* variants across all *N* tumors for a fixed value of index *b*.

## Results

### Proportionate Contributions of Mutational Processes to Cancer Effect Can Be Calculated

To determine the sources of mutagenesis occurring in tumor samples, we deconvolved the mutational burden of each tumor into the most likely distribution of attributed single-nucleotide variant mutational signatures (Alexandrov and Zhivagui, 2018; Petljak and Alexandrov, 2016). Applied to a lung squamous cell carcinoma (LUSC) tumor sample from a single patient (MDA-1229-T), this deconvolution yielded three trinucleotide mutational signatures (age-associated #5, tobacco smoking #4, and unknown aetiology #8; **Fig. 1A**), each contributing to the flux of single nucleotide variants in the tumor at a calculated weight (**Fig. 1B**). The trinucleotide signature weight or combination of trinucleotide signature weights contributing to a specific variant (**Fig. 1A**) times the proportion of mutational causation attributable to each corresponding cancer effect (**Fig. 1B**) provides the probability each source contributed to each variant in this tumor (**Fig. 1C**). In this instance, age-associated signature #5 is the most likely contributor to TP53 R282W and NFE2L2 R34P, whereas tobacco smoking is the most likely source of mutations causing OR13G1 L79L. However, only a few of the mutations that occur in somatic tissue are thought to be selected for their effects on growth or survival, and therefore causative of cancer, and the level of causation is presumably quantitative—i.e., the mutations in a type of cancer that drive cancer are responsible to different degrees for the manifestation of a cancer phenotype (Cannataro et al., 2018). In this case, TP53 R282W has a higher cancer effect size than NFE2L2 R34P, and the odorant receptor mutation OR13G1 L79L has negligible to no effect. The product of the probability that each mutational source contributed to each variant in this tumor and the effect of the specific variant (**Fig. 1D**) quantifies the probability-weighted cancer effect for each variant by each source (**Fig. 1E**). Summing the probability-weighted cancer effect for each source across variants yields the proportion of cancer effect attributable to each source of mutations (**Fig. 1F**). Age-associated mutational signature #5 contributed the highest weight in MDA-1229-T, and led to the largest estimated effect through its high probability of being causative of both the NFE2L2 R34P and TP53 R282W mutations. Via deconvolution of the mutational signatures responsible for recurrent variants in cancer and calculation of the cancer effect sizes of the nucleotide substitutions driving cancer evolution, we have calculated which mutagenic sources fueling nucleotide variation can be attributed as proportionally causative of individual tumors in patients.

### Mutagenic Input and Cancer Effect from each Source Can Differ Substantially within Tumors

The match between the proportional input to total mutations by each mutagenic source (**Fig. 1B**) and the proportional cancer effect arising from each mutagenic source (**Fig. 1F**) varies in each patient’s tumor (**Fig. 2**). Quantifying the match by the Jensen-Shannon Divergence (JSD) between proportional mutational input and proportion of cancer causation, we found the tumor type with the lowest median divergence to be ovarian serous cystadenocarcinoma (OV, **Fig. 2A**). The mutational input to the OV tumor in **Fig. 2A** with the lowest JSD (TCGA-24-1103) was entirely attributed to the BRCA-1- and BRCA-2-associated signature (#3); thus, all cancer effects were attributable to this single source of mutation. Indeed, tumor sample TCGA-24-1103 has a somatic BRCA2 L1638E mutation. Examining a tumor at the second quartile of the distribution of JSD (TCGA-09-0366; **Fig. 2A**), there is a slight mismatch between the mutational input and the contribution to cancer causation—with the clock-like signature #5 exhibiting slightly more cancer effect than signature weight. Three additional tumors drawn at the median JSD, the fourth quintile, and the highest JSD demonstrate increasing degrees of mismatch between the mutational input and the contribution to cancer causation. These mismatches are even more frequent in 22 other tumor types analyzed (rectal adenocarcinoma, at approximately the second quartile of median JSD across tumor types, **Fig. 2B**; human papillomavirus virus negative head and neck squamous-cell carcinoma, at approximately the third quartile of median JSD across tumor types, **Fig. 2C**; and low-grade glioma, exhibiting the greatest median mismatch across tumor types, **Fig. 2D**; **Supplemental Fig. 2**).

**Figure 2.**
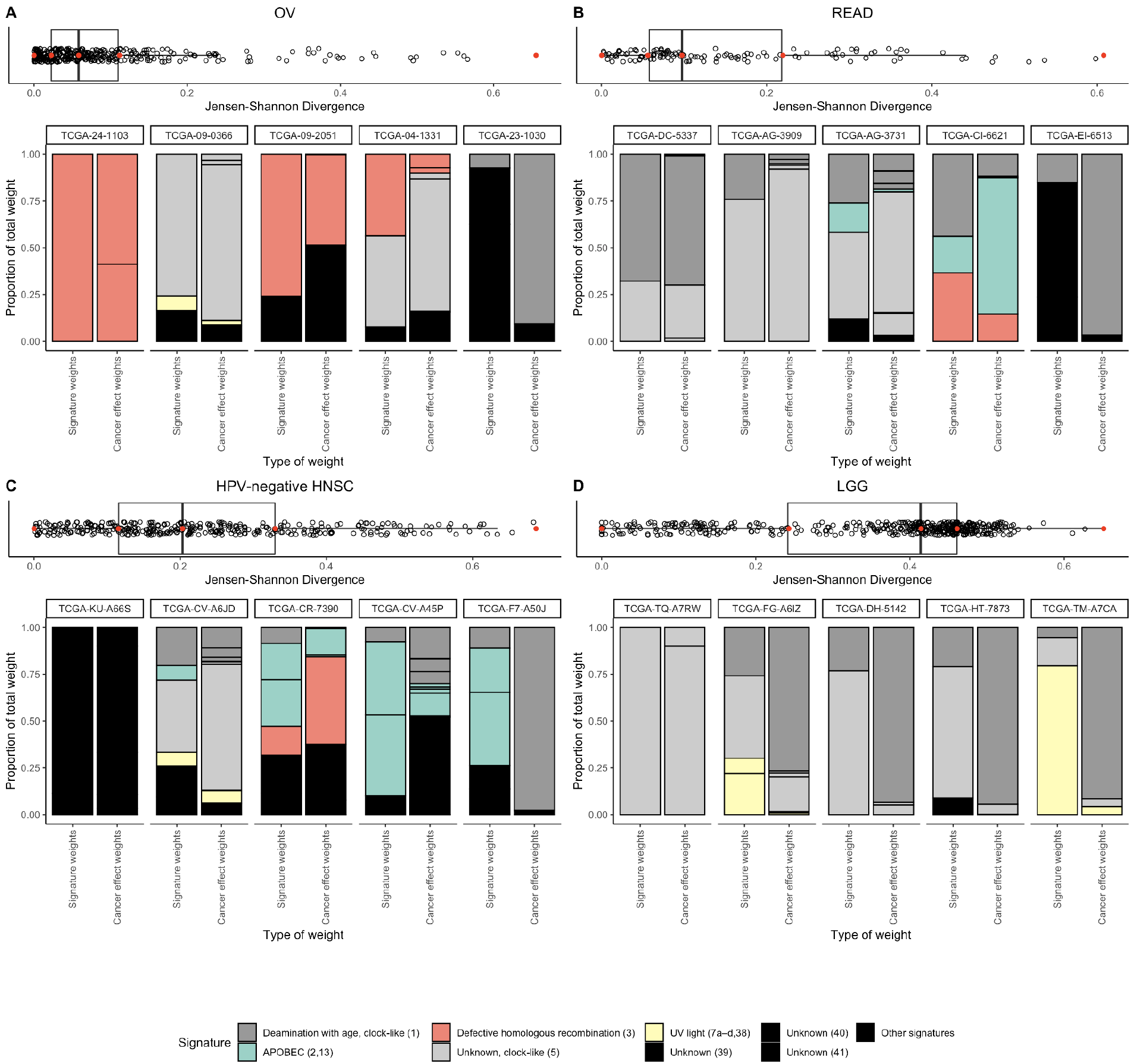
Box plots of the Jensen-Shannon Divergence (JSD) between the proportional input of each mutagenic source and the proportion that each signature contributed to the total cancer effect in each tumor, accompanied by the proportions of observed mutations in a tumor caused by each signature and the proportions of cancer-causation attributable to each source of mutations, for the five tumors (red dots) bounding the quartiles of the JSD among tumors, for four cancer types that bound the tertiles of median JSD among cancer types: (**A**) the cancer type with the least median divergence between the proportional input of each mutagenic source and the proportion that each signature contributed to the total cancer effect, ovarian serous cystadenocarcinoma (OV), (**B**) rectum adenocarcinoma (READ), (**C**) HPV-negative head-and-neck squamous-cell carcinoma (HPV-negative HNSC), and (**D**) the cancer type with the greatest median divergence between the proportional input of each mutagenic source and the proportion that each signature contributed to the total cancer effect, lower-grade glioma (LGG).

### Mutagenic Input and Cancer Effect from each Source Can Differ Enormously among Oncogenic Variants within each Cancer Type

Many well-known processes have been established as major contributors to tumor mutation burden, such as tobacco in lung tissues, ultraviolet radiation in skin tissues, and APOBEC cytidine deaminases in bladder, cervical, and HNSC tissues. However, mutational processes are trinucleotide-specific, which leads to differences in underlying amino-acid mutation rates depending on the sequence context of each variant site. The mutational process mostly likely to originate an oncogenic variant can not only differ from variant to variant, but can also differ from the mutational process that causes the greatest number of mutations within each tumor type (**Fig. 3**). For instance, among actionable processes, mutations in lung adenocarcinoma and lung squamous-cell carcinoma were most frequently attributed to tobacco-associated mutagenesis (**Fig. 3A–B**). The high attribution of KRAS G12C mutations to this lung-specific mutagenic process explains their high frequency in LUAD compared to other RAS-driven cancer types such as pancreas or colon adenocarcinomas. Major driver variants of KRAS and TP53, in LUAD and LUSC respectively, exhibit markedly different origination rates from tobacco-associated processes. Perhaps most notable is the minimal attribution of EGFR L858R to tobacco-associated mutagenic processes. The attribution of tobacco-associated mutagenic processes to the cancer effects of KRAS G12 variants and EGFR L858R (**Fig. 3A–B**) are consistent with—and provide an explanation for—the increased odds of KRAS mutation in tumor tissue of ever smokers compared to never smokers, as well as the increased odds of EGFR mutation in never smokers compared to ever smokers (Chapman et al., 2016). Even nucleotide variants that do not cause an amino-acid substitution have quantifiable cancer effects that can be attributed to mutagenic processes—e.g. TP53 T125T, which affects splicing of the TP53 transcript (Varley et al., 2001), is attributable to tobacco in both LUSC and LUAD; **Fig. 3A–B**(Varley et al., 2001). Ultraviolet light (UV) is the major mutagenic process leading to both total mutations and most major oncogenic variants in primary skin cutaneous carcinoma (SKCM, **Fig. 3C**). SKCM oncogenic variants are dominated by the high effect size of UV-driven BRAFV600E (cf. Cannataro et al., 2018), but one major oncogenic variant common to SKCM (KIT K642E) is almost entirely attributable to age-associated processes rather than UV (**Fig. 3C**). Many of the high-effect mutations of CTNNB1 in LIHC are attributable to mutational processes generating (COSMIC Signature 16; Letouzé et al., 2017) (**Fig. 3D**). The greatest proportion of cancer effect for several oncogenic somatic variants in LIHC—such as TP53 R249S and CTNNB1 D32V—is attributable to mutagenic chemical exposure; and the greatest proportion of cancer effect in several other CTNNB1 variants are attributable to processes with as-yet unknown etiology that may in the future be linked to other mutagenic chemical exposures. Mutations in bladder urothelial carcinoma (BLCA) were most frequently attributed to APOBEC cytidine deaminases that are thought to be activated by exposure to viruses, which may be presumed to be preventable. However, 7 of the top 10 variants as determined by cancer effect were attributed to non-actionable, age-associated processes rather than to APOBEC-associated mutagenic processes. In contrast, three known cancer driver variants (FGFR3 S249C, PIK3CA E545K, and PIK3CA E542K), were almost entirely attributed to the action of APOBEC cytidine deaminases. Cervical squamous-cell carcinoma and endocervical adenocarcinoma (CESC), human-papillomavirus-negative head-and-neck squamous-cell carcinoma (HNSC HPV negative), and human-papillomavirus-positive head-and-neck squamous-cell carcinoma (HNSC HPV positive) were also dominated by APOBEC-associated mutations. CESC and HNSC also exhibited diversity in which process was most likely to originate each oncogenic variant (Cannataro et al., 2019); however, attributions of APOBEC-associated processes for the origination of oncogenic PIK3CA E542K and PIK3CA E545K mutations are consistent across multiple cancer types (cf. **Fig. 3B, E–H**).

**Figure 3.**
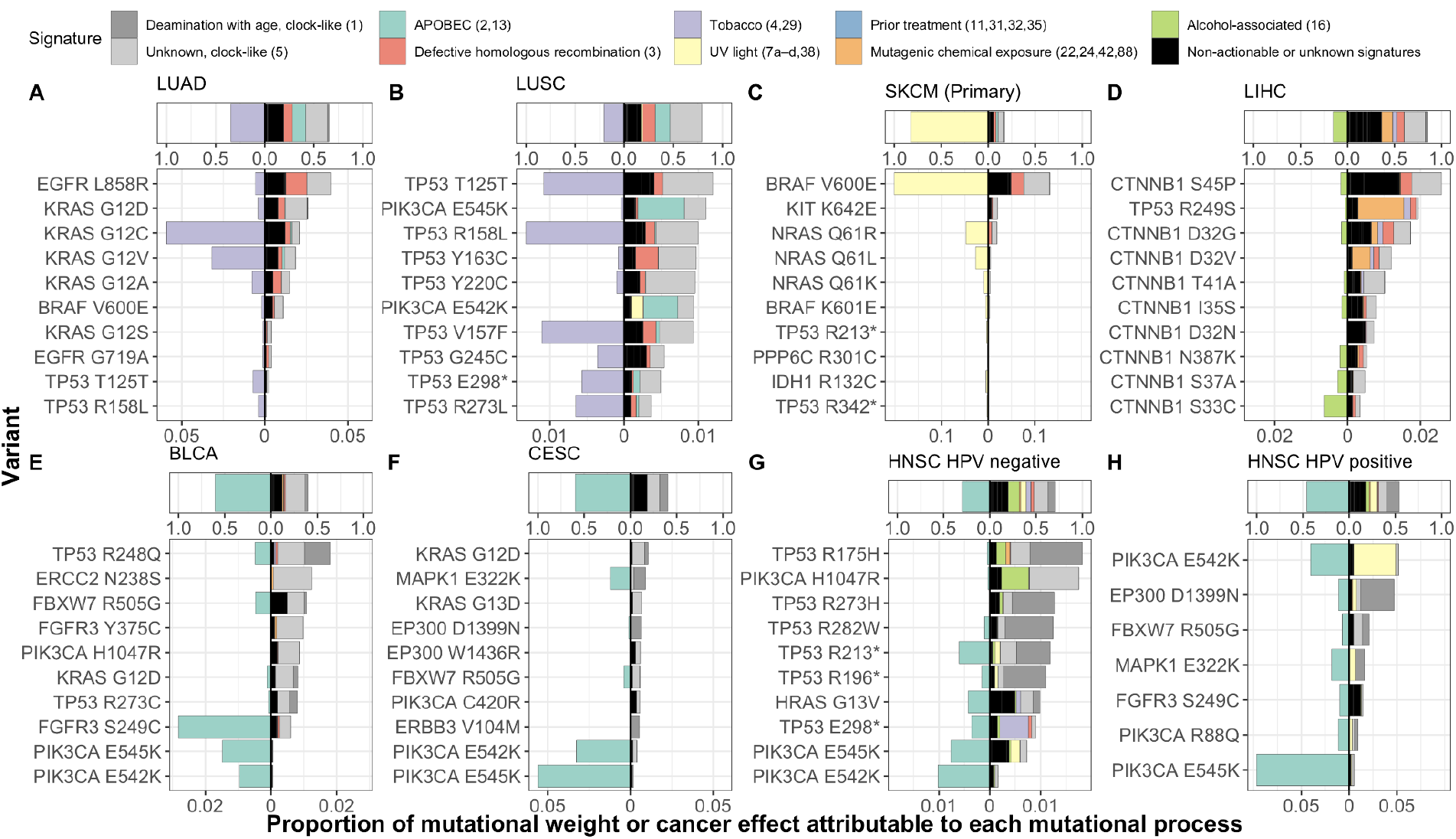
In each cancer type, the contribution of each mutational process to total mutation burden (top) and to the average cancer effect of variants classified as drivers (bottom). Cancer effect is quantified proportionate to the total cancer effect in each tumor, and variants are filtered by gene drivers defined in (Bailey et al., 2018)). For each cancer type, the average contribution of the dominant actionable process (quantified by the bar width left of the *x*-axis origin) and the non-dominant or non-actionable processes (quantified by the bar width right of the *x*-axis origin) to total mutation is shown, above the top 10 variants contributing the greatest cancer effect to primary tumors of that cancer type, ordered by the average proportional effect size attributable to that variant from all of the non-dominant or non-actionable mutational processes, alongside the average contribution of the dominant actionable processes (left of the *x*-axis origin) and the non-dominant or non-actionable processes (right of the *x*-axis origin) to cancer effect (measured proportionate to the total effect in each tumor). (**A**) Lung adenocarcinoma (LUAD), (**B**) Lung squamous-cell carcinoma (LUSC), (**C**) Skin cutaneous melanoma (SKCM, primary tumors only), (**D**) Liver hepatocellular carcinoma (LIHC), (**E**) Bladder urothelial carcinoma (BLCA), (**F**) Cervical squamous-cell carcinoma and endocervical adenocarcinoma (CESC), (**G**) Human-papillomavirus-negative head-and-neck squamous-cell carcinoma (HNSC HPV negative), (**H**) Human-papillomavirus-positive head-and-neck squamous-cell carcinoma (HNSC HPV positive).

### Relative Mutagenic Input and Relative Cancer Effect are Specific to each Tumor Type

The mismatches between the proportional input to total mutations by each mutagenic source (**Fig. 1B**) and the proportional cancer effect arising from each mutagenic source (**Fig. 1F**) exist not only at the level of individual tumors, but also at the level of tumor types—where they indicate which mutational sources make an outsized contribution to the causation of cancer compared to their production of mutations, and *vice versa*. Many tumor-type mutational-signature pairs exhibit statistically significant differences between the proportional input to total mutations by each mutagenic source and the proportional cancer effect arising from each mutagenic source (continuity-corrected Wilcoxon two-sided rank-sum tests, *P* < 0.05; **Fig. 4A**). For example, APOBEC-related signatures 2 and 13 exhibit larger mutation weight than cancer effect across many cancer types, as do 17a, 21, 22, 26, 28, and 37. In contrast, the aging-associated signature 1 exhibits larger cancer effect than mutation weight across many cancer types, as does polymerase-epsilon signature 10b. In lower-grade glioma, the age-associated signature 5 constitutes much more of the mutation weight than its cancer effect (68% compared to 23%), whereas age-associated signature 1 has the opposite relationship (23% compared to 78%). This difference between the two age-associated signatures is largely attributable to the high effect size of IDH1 variants, which occur predominantly as a consequence of ACG→ATG mutations that are frequent in signature 1 and rare in signature 5. A similar contrast can be seen in thyroid adenocarcinoma, wherein the APOBEC-related signature 2 exhibits high mutation weight and virtually zero cancer effect (30% compared to 0.2%), and wherein the aging signature 5 exhibits much more cancer effect than mutation weight (68% compared to 30%). This contrast comes about because thyroid adenocarcinoma is often driven by BRAF V600E mutations that convey enormous cancer effects, and BRAF V600E mutations come about frequently as a consequence of GTG→GAG mutations that are found at low frequency within the aging signature 5, but are found at extremely low frequency within APOBEC signature 2.

**Figure 4.**
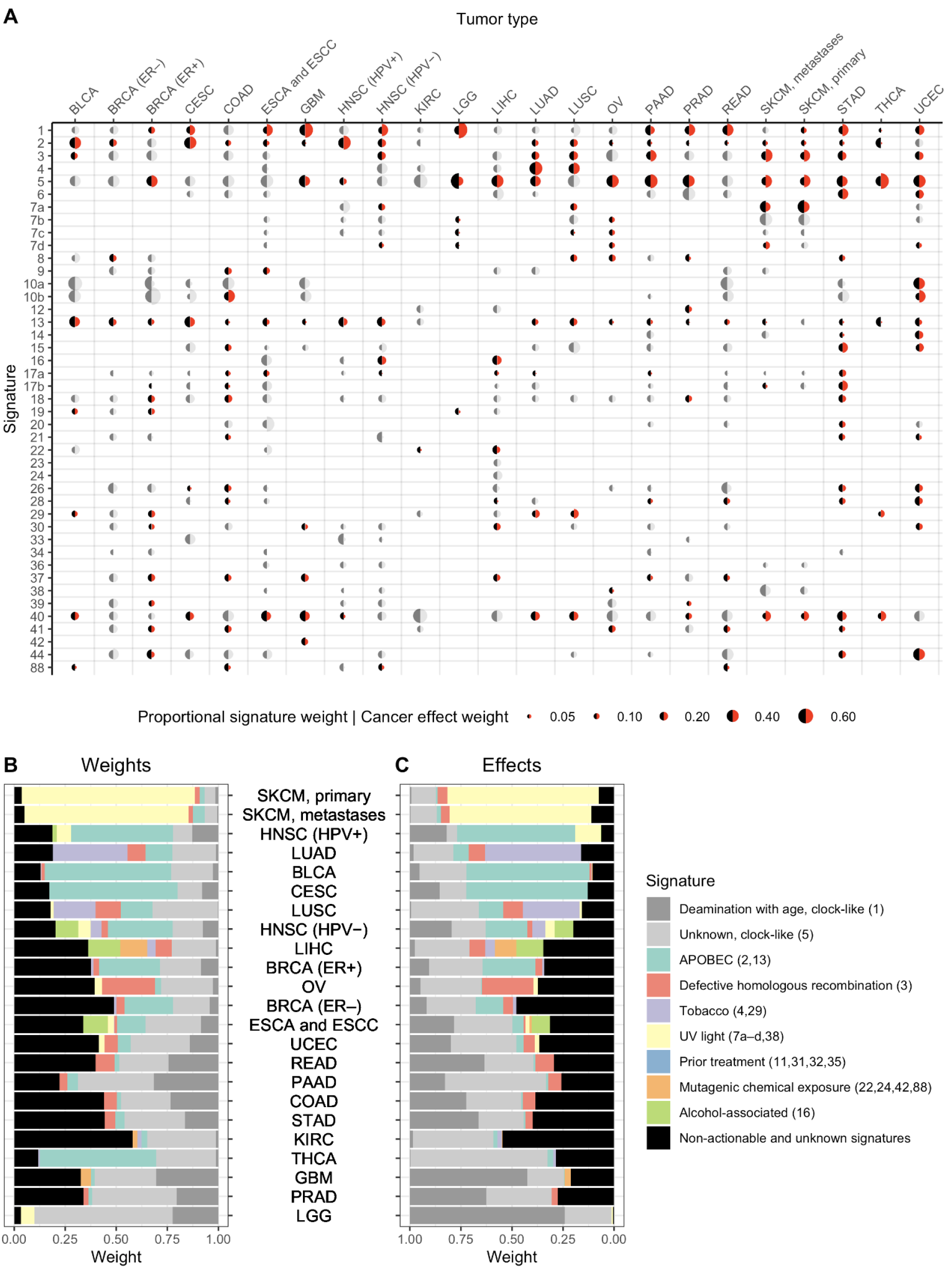
Mutational process weights and cancer effects for unknown and etiology-associated mutation signatures. (**A**) Relative contributions of processes attributed to each mutational signature across 23 cancer types to total SNVs (black half-circle) and cancer effects (red half-circle; effects that are not statistically significantly different by a paired Wilcoxon two-sided rank-sum test are indicated in dark and light gray half-circles). For each cancer type, some mutagenic processes contribute lesser proportions of cancer effect than the proportion of SNVs that they cause (e.g. SBS1 in PAAD; 5 in LGG; 7a and 7b in SKCM; 13 and 40 in THCA), and some mutagenic processes contribute greater proportions of cancer effect than the proportion of SNVs that they cause (e.g. SBS1 in ESCA, HNSC, LGA, PRAD, and STAD; 2 in HPV^+^ HNSC; 5 and 40 in THCA). (**B**) Relative contributions of age-associated (grey), actionable (colors), and unknown processes (black) across 23 cancer types to total SNVs. (**C**) Relative cancer effects of age-associated, actionable, and unknown processes.

### Preventable Mutational Processes Contribute Substantially to Causation of Skin, Head-and-Neck, Cervical, and Lung Cancer

Among the non-age-related etiologies are a number of mutational processes that are putatively “preventable”—in that they can be mitigated by individual behaviors or interventions (**Fig. 4B–C**). Skin cancer, lung cancer, HPV-positive head and neck cancer, and cervical cancer are notable for the dominant role of putatively preventable processes underlying both raw SNV mutation weight (**Fig. 4A**) and cancer effect (**Fig. 4B**). Lower-grade glioma, glioblastoma, and prostate adenocarcinoma are notable for the lack of putatively preventable processes underlying both raw mutation weight and cancer effect. However, the mutation weights and cancer effects are not the same. For example, 57% of the mutation weight of thyroid adenocarcinoma is associated with APOBEC processes that might be preventable by avoiding viral infections. However, these mutations contribute only 2.9% of the cancer effect (*P* < 0.001, Wilcoxon rank-sum test), so that preventing APOBEC-associated mutation would likely do little to prevent the majority of THCA cancer. A contrasting case is the lung cancers: the net cancer effects of the SNVs attributable to tobacco chewing (3% in LUAD and 4% in LUSC) and tobacco smoking (44% in LUAD and 24% in LUSC) are larger than the mutation weights of these sources of mutagenesis (2% and 2% for tobacco chewing and 35% and 19% for smoking in LUAD and LUSC, respectively; *P* < 0.001 for signature 4 for both lung cancer types and for signature 29 in LUSC; *P* = 0.008 for signature 29 in LUAD; Wilcoxon rank-sum test).

### Age-associated Mutational Processes Contribute Substantially to Causation of Glioma, Prostate, Thyroid, Pancreatic, and Colorectal cancer

Because each mutagenic process is linked to a trinucleotide variant signature that has been identified as clock-like (ubiquitous and age-associated) or non-clocklike (Alexandrov et al., 2020, not associated with age; 2015), the proportion of total mutations attributable to clock-like processes and non-clocklike processes in each cancer type can be quantified (**Fig. 4B**). Among tissues, the cancer types with the greatest proportion of total mutations contributed by non-clocklike processes are melanoma (primary and metastatic), cervical squamous cell carcinoma and endocervical adenocarcinoma, and head and neck cancers. Lower grade glioma exhibits the greatest proportion of total mutations contributed by age-associated, “clocklike” processes. Moreover, with regard to the explanation of tumorigenesis and cancer incidence, the cancer effect attributable to age-associated processes and non-age-associated processes in each cancer type can be quantified (**Fig. 4C**). Among tumor tissues, those with the greatest proportion of cancer effect contributed by age-associated processes are gliomas (LGG, GBM) and prostate adenocarcinomas, consistent with the strong association of the incidence of these cancers with age (Dubrow and Darefsky, 2011; Rawla, 2019), as well as pancreatic cancers. Primary and metastatic melanoma, lung adenocarcinoma, and HPV-positive head and neck squamous-cell carcinoma exhibit the greatest proportion of cancer effect contributed by non-age-associated processes, consistent with the strong association of these cancers with exogenous factors (UV exposure, smoking, and HPV infection).

Our analysis attributes an amount of cancer causation to such endogenous and inactionable processes that varies widely among cancer types. Cancer types varied in the degree to which their causation was associated with COSMIC signature #1, which correlates with stem cell division in different tissues and represents the processes associated with the mitotic clock (Tomasetti et al., 2017; Fig. 3C; c.f. Tomasetti and Vogelstein, 2015), ranging from extremely small contributions (<0.05%) in THCA, LUSC, SKCM, KIRC, LUAD, LIHC, and BLCA, to 76% of the cancer effect in LGG. Combining the replication-associated and non-replication-associated aging signatures, causation attributable to all aging-associated processes ranged from 13% in SKCM to 99% in LGG; in around half of the cancer types a minority of cancer causation was attributable to age-associated processes. Age has not been associated across cancer types with any of the signatures that have unknown etiology. Greater than 50% of the cancer effect leading to KIRC is attributed to unknown mutational processes; OV, PRAD, and BRCA (ER-) all have >30% of their cancer effect caused by currently unknown or unattributed mutational processes (**Fig. 4C**).

## Discussion

Here we have shown that the impact on carcinogenesis of mutagenic processes associated with single-nucleotide variant signatures can be quantified. This quantification is distinct from the number or proportion of mutations that can be attributed to a process, because it accounts for the extent to which each mutation contributes to the cancer phenotype—increased replicative and survival advantage in each tissue and cancer type—via single-nucleotide variants. We have shown how to use the proportions of observed mutations in a tumor caused by each signature to calculate the probability that each mutational source contributed to each variant in this tumor. Each of these probabilities serves to weight the cancer effect size of each variant, yielding the probability-weighted portion of effect size for each variant attributable to each source of mutations and thus the proportion of cancer-causation attributable to each source of mutations. In turn, the quantification of cancer-causation within each tumor, characterized across a population of patients, provides a reductionist molecular approach toward quantifying the degree to which a process can be held responsible for carcinogenesis in a cancer type that is wholly distinct from traditional epidemiological studies.

Our analysis of the cancer effects of single-nucleotide mutations and associated signatures has been enabled by quantitative estimates of their intrinsic mutation rates (Fousteri and Mullenders, 2008; Lawrence et al., 2013; Stamatoyannopoulos et al., 2009). Deconvolution of the quantitative contributions of known mutation signatures explains the high prevalence of KRAS G12C and low prevalence of EGFR L858R in ever-smokers, and the converse relationships in never-smokers. It illuminates the potent role of ultraviolet light in BRAF V600E-driven melanoma. It attributes major drivers PIK3CA E542K and E545K to the potentially virally-induced action of APOBEC cytidine deaminases, and highlights unknown processes that deserve further identification such as those underlying high-cancer-effect single-nucleotide variants of LIHC. Importantly, germline variants, copy-number variation, epigenetic alterations, and changes to the aging tissue microenvironment also contribute to the cancer phenotype (Laconi et al., 2020; Liggett and DeGregori, 2017; Montgomery et al., 2018; Mroz et al., 2015; Ramakodi et al., 2016; Sun et al., 2018). Incorporation of signatures associated with these kinds of alterations (Macintyre et al., 2018) and of attributions of each signature to relevant sources would markedly increase the purview of inferred cancer causation, revealing a full picture of the importance of diverse mechanisms behind the spectrum of genomic alterations fueling cancer evolution.

For an individual cancer patient, calculation of the relative cancer effect of diverse sources of mutation provides an estimate of how much each mutagenic process is responsible for an individual’s cancer. From a public health perspective, these calculations constitute a bridge between molecular studies and long-standing epidemiological analyses that have associated behaviors (e.g., smoking) or professions (e.g., sun exposure) with cancer incidence. Public health intervention targeted at minimizing exposure to these actionable signatures would mitigate disease severity by preventing the accumulation of mutations that directly contribute to the cancer phenotype. Finally, our findings connect specific mutagenesis patterns and processes with cancer, providing a “smoking gun” that can inform individuals as to why an instance of cancer happened—and have promise to play a significant role in demonstrating individual as well as group-level cause for legal recourse due to carcinogenic exposure (e.g. Lee, 2016).

The quantification of cancer effect attributable to specific sources of mutation has evident parallels to epidemiological results that assess the effect of risk factors on cancer causation (Shield et al., 2016). These epidemiological results often rely on correlation, and calculate an increase in the probability of cancer in relation to some behavior or exposure. Calculations of the relative cancer effect of diverse sources of mutation, in principle, directly relates the mutations driving tumorigenesis to mechanistic processes. However, multiple challenges impede their use at a population level in comparison to longstanding, well-crafted epidemiological studies: 1) conducting appropriate tumor sampling—most large tumor sequencing studies are sampled haphazardly, without reference to a distinct population, without stratification or even “random” sampling; 2) formulating an “apples to apples” quantitative mapping comparing proportions of effect to odds ratios; and 3) forming a discrete mapping of mutational signatures to mechanistic processes to epidemiological factors. These attributions of cause associated with COSMIC signatures are critical to our interpretations of these results, and range in surety from well-established (e.g., UV #7 and smoking #4), to presumptive (e.g. indirect damage from UV light #38).

Recent research has touched on a debate as to what extent “bad luck”—endogenous mutagenic processes that accumulate naturally with age—plays a role in the incidence of cancer arising in various tissues. Here, we addressed the question regarding the relative contributions of exogenous and endogenous sources of mutation to tumorigenesis by quantifying the extent that specific variants are driving tumorigenesis, and attributing the variants back to the mutational processes that originally fueled their creation. We found that signatures relating to aging processes (#1 and #5) were responsible for the majority of cancer effect in tumors of the brain (LGG, GBM) and tissues with large amounts of epithelial turnover (READ, COAD, STAD, UCEC). Other tumors whose cancer effects could largely be attributed to aging include PRAD—a tumor type strongly associated with age (Bostwick et al., 2004), THCA (whose major single-nucleotide driver, BRAFV600E, is more likely to be caused by mutations associated with clock-like signature #5 than by mutations associated with other signatures), and PAAD. Several tumor types have large proportions of the cancer effect size directly attributable to mutational processes that are actionable, i.e., interventions could reduce the mutations in these tissues that are responsible for the cancer-causing variants. CESC, HNSC, and BLCA are largely driven by mutations attributed to virus-induced APOBEC activity, SKCM is largely driven by UV light exposure, and mutations responsible for increased proliferation and survival of cancerous cells within lung cancers trace back to smoking.

The importance of understanding the underlying sources of mutations that ultimately lead to cancer in each and every patient—whether they are endogenous or exogenous, and whether they come from sources that are actionable—is underscored by the remarkable successes of anti-smoking interventions against carcinogenic exposures, which have saved many lives (Holford et al., 2014). In our study, some cancer types such as KIRC, BRCA (ER-), OV, and PRAD exhibited a large proportion of cancer effect that was attributable to signatures with unknown etiology. As we gain greater insight into these mutational signatures and their diverse causative mechanisms, we may discover additional actionable mutational processes that can be mitigated by proactive public health interventions.

## Acknowledgements

Members of the Townsend Lab provided stimulating discussions and helpful feedback on this research. Funding for this research was provided by NIH 1P50DE030707, NIH 1R01LM012487, NIH 1R01CA215900, NIH 5R01 CA231112, the Yale Cancer Biology Training program (NIH T32 CA193200/CA/NCI HHS/United States), and the Elihu Professorship endowed research funds.

## Author Contributions

VLC and JPT designed the research. VLC assembled all data and executed all analyses. JDM provided computational tools and expertise. VLC and JPT wrote the manuscript. All authors reviewed the manuscript.

---

**Supplemental Table S1:**
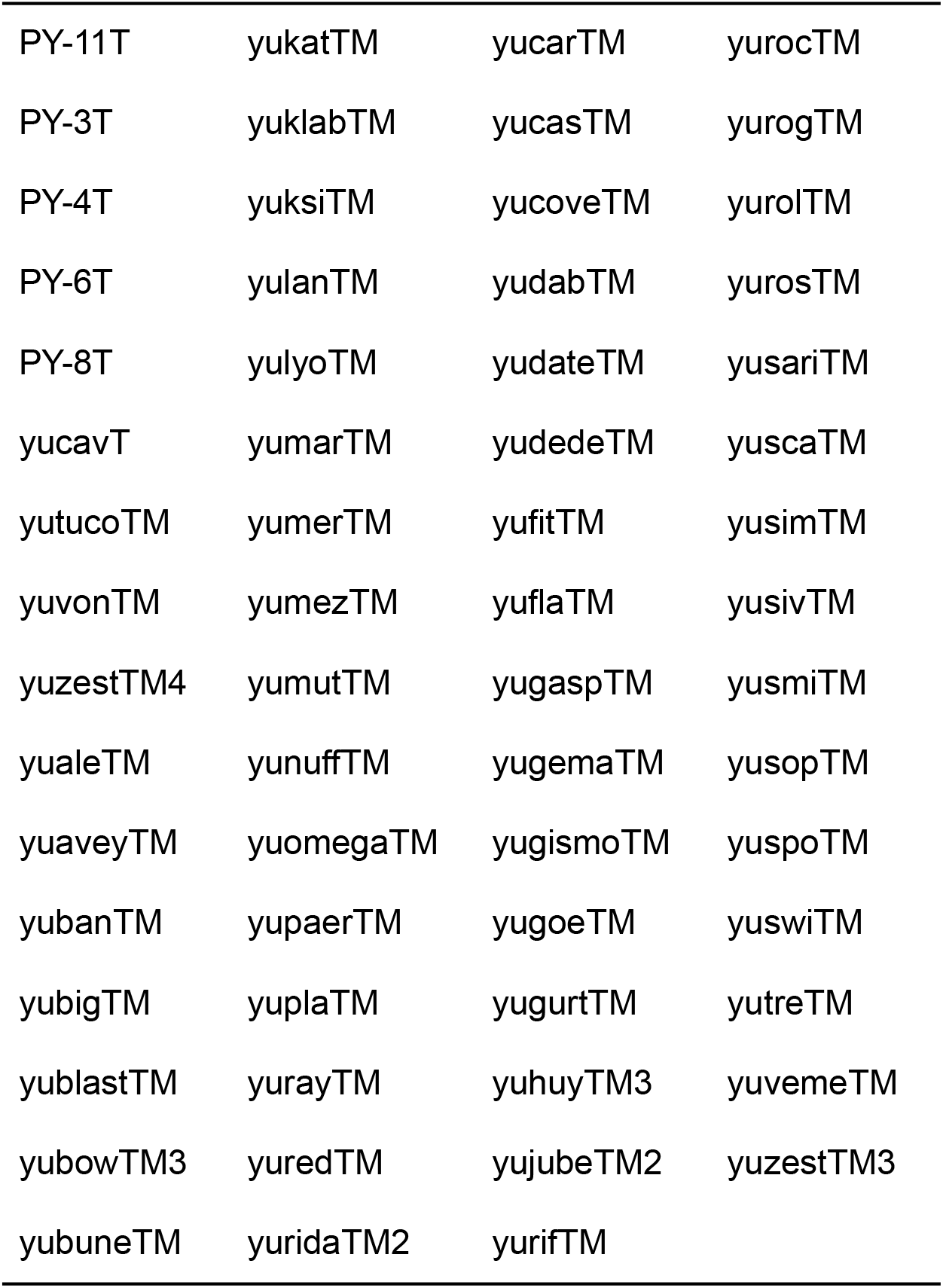
Yale-Gilead sequenced tumors that received chemotherapies and were removed from this analysis

**Supplementary Figure S1:**
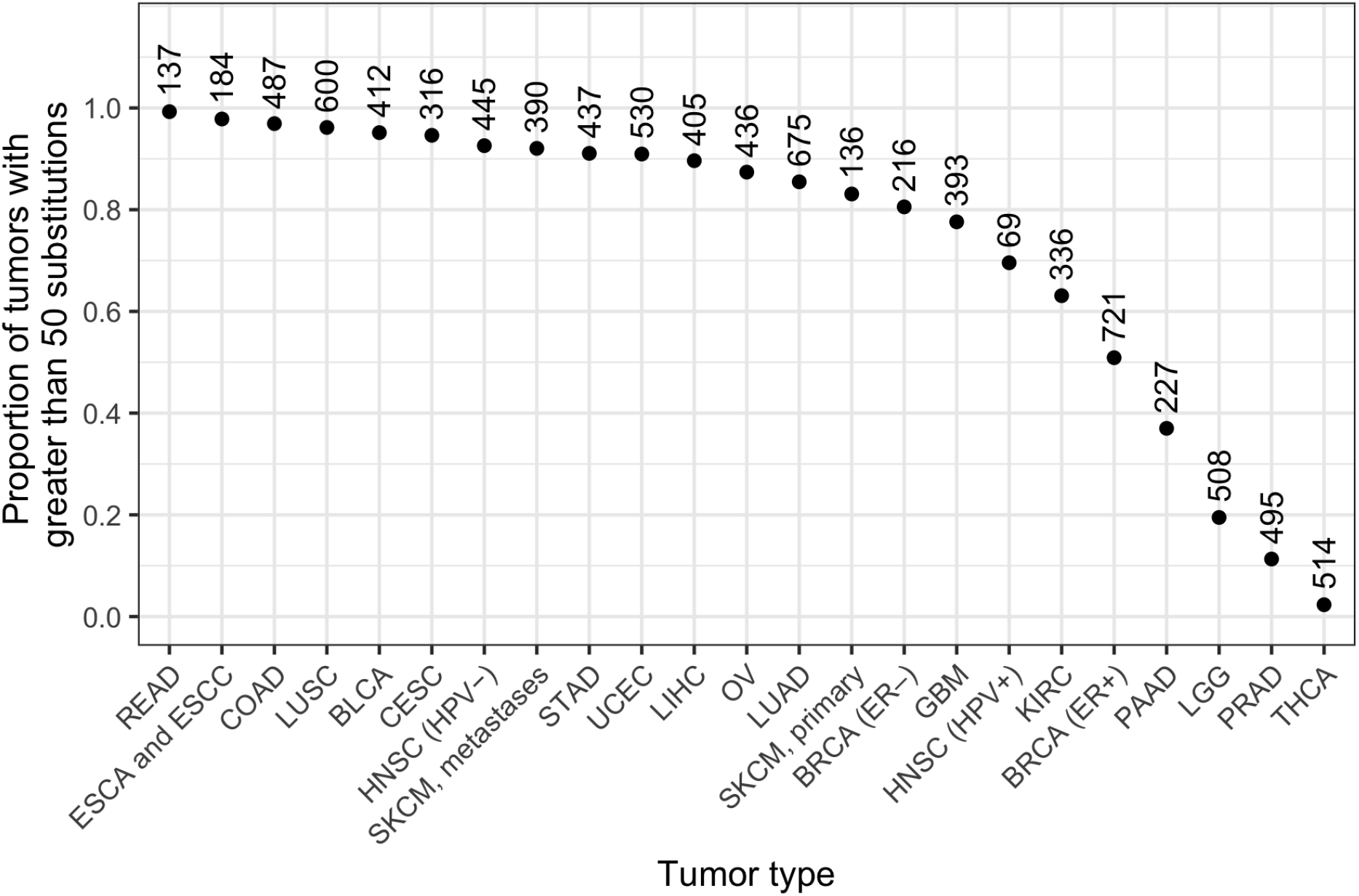
The proportions of tumors with greater than 50 substitutions in 23 tumor types. Numbers above the points indicate the total number of tumors in the data subset.

**Supplemental Figure S2.**
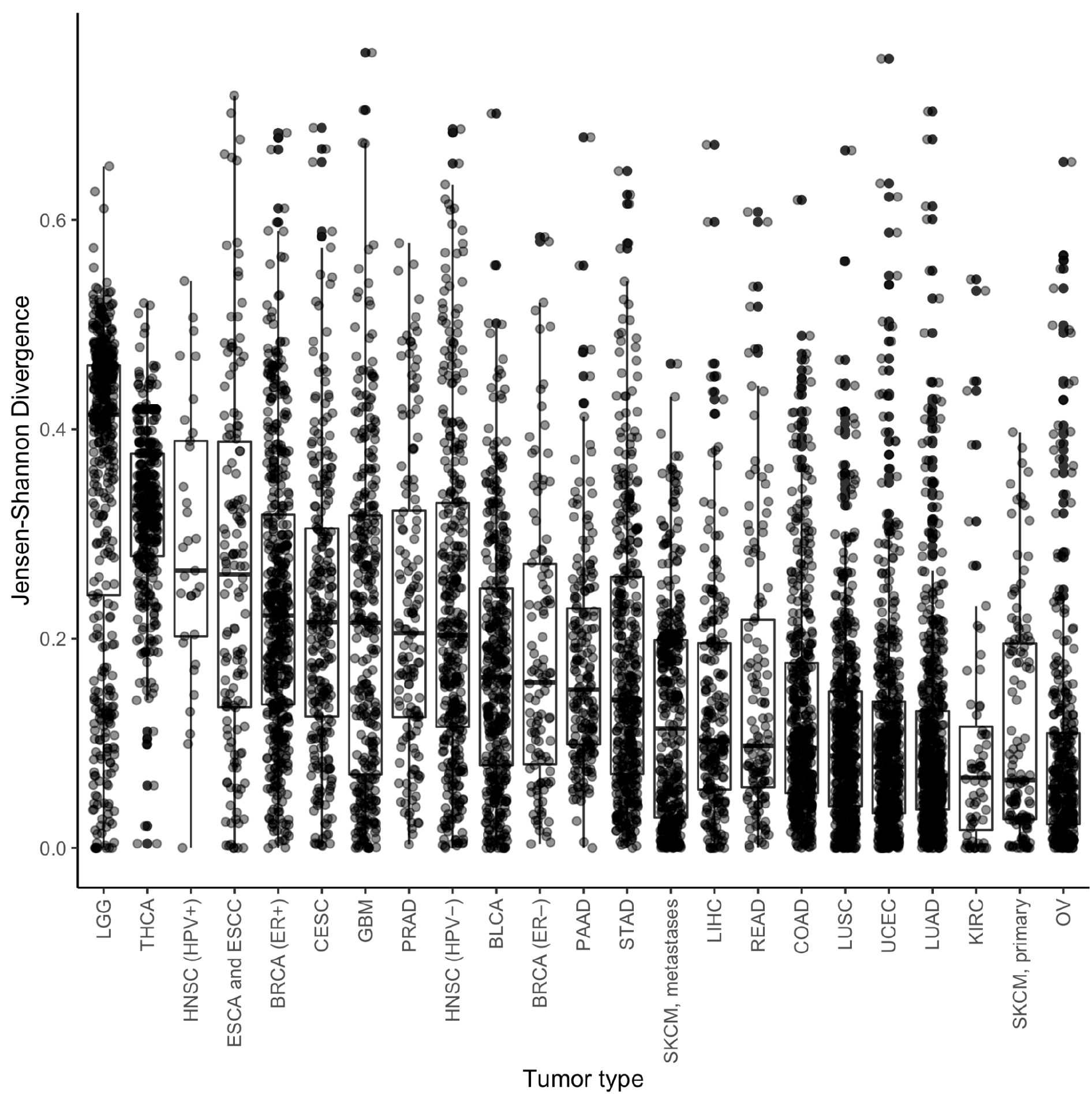
Box plots by tumor type of the Jensen-Shannon Divergence between the proportional input of each mutagenic source and the proportion that each signature contributed to the total cancer effect in each tumor, for 23 cancer types: brain lower-grade glioma, thyroid carcinoma, HPV^+^ head and neck squamous-cell carcinoma, esophageal cancers, ER^+^ breast invasive carcinoma, cervical squamous cell carcinoma and endocervical adenocarcinoma, glioblastoma multiforme, prostate adenocarcinoma, HPV^-^ head and neck squamous-cell carcinoma, bladder urothelial carcinoma, ER^-^ breast invasive carcinoma, pancreatic adenocarcinoma, stomach adenocarcinoma, skin cutaneous melanoma (metastatic), liver hepatocellular carcinoma, rectum adenocarcinoma, colon adenocarcinoma, lung squamous-cell carcinoma, uterine corpus endometrial carcinoma, lung adenocarcinoma, kidney renal clear cell carcinoma, skin cutaneous melanoma (primary), ovarian serous cystadenocarcinoma.

## References cited

Alexandrov LB, Jones PH, Wedge DC, Sale JE, Campbell PJ, Nik-Zainal S, Stratton MR. 2015. Clock-like mutational processes in human somatic cells. Nat Genet 47:1402–1407.

Alexandrov LB, Kim J, Haradhvala NJ, Huang MN, Tian Ng AW, Wu Y, Boot A, Covington KR, Gordenin DA, Bergstrom EN, Islam SMA, Lopez-Bigas N, Klimczak LJ, McPherson JR, Morganella S, Sabarinathan R, Wheeler DA, Mustonen V, PCAWG Mutational Signatures Working Group, Getz G, Rozen SG, Stratton MR, PCAWG Consortium. 2020. The repertoire of mutational signatures in human cancer. Nature 578:94–101.

Alexandrov LB, Zhivagui M. 2018. Mutational Signatures and the Etiology of Human Cancers. Reference Module in Biomedical Sciences. doi:10.1016/b978-0-12-801238-3.65046-8

Ashford NA, Bauman P, Brown HS, Clapp RW, Finkel AM, Gee D, Hattis DB, Martuzzi M, Sasco AJ, Sass JB. 2015. Cancer risk: Role of environment. Science 347:727–727.

Bailey MH, Tokheim C, Porta-Pardo E, Sengupta S, Bertrand D, Weerasinghe A, Colaprico A, Wendl MC, Kim J, Reardon B, Kwok-Shing Ng P, Jeong KJ, Cao S, Wang Z, Gao J, Gao Q, Wang F, Liu EM, Mularoni L, Rubio-Perez C, Nagarajan N, Cortés-Ciriano I, Zhou DC, Liang W-W, Hess JM, Yellapantula VD, Tamborero D, Gonzalez-Perez A, Suphavilai C, Ko JY, Khurana E, Park PJ, Van Allen EM, Liang H, MC3 Working Group, Cancer Genome Atlas Research Network, Lawrence MS, Godzik A, Lopez-Bigas N, Stuart J, Wheeler D, Getz G, Chen K, Lazar AJ, Mills GB, Karchin R, Ding L. 2018. Comprehensive Characterization of Cancer Driver Genes and Mutations. Cell 174:1034–1035.

Barnes JL, Zubair M, John K, Poirier MC, Martin FL. 2018. Carcinogens and DNA damage. Biochem Soc Trans 46:1213–1224.

Bostwick DG, Burke HB, Djakiew D, Euling S, Ho S-M, Landolph J, Morrison H, Sonawane B, Shifflett T, Waters DJ, Timms B. 2004. Human prostate cancer risk factors. Cancer 101:2371–2490.

Brash DE, Rudolph JA, Simon JA, Lin A, McKenna GJ, Baden HP, Halperin AJ, Pontén J. 1991. A role for sunlight in skin cancer: UV-induced p53 mutations in squamous cell carcinoma. Proc Natl Acad Sci U S A 88:10124–10128.

Cannataro VL, Gaffney SG, Sasaki T, Issaeva N, Grewal NKS, Grandis JR, Yarbrough WG, Burtness B, Anderson KS, Townsend JP. 2019. APOBEC-induced mutations and their cancer effect size in head and neck squamous cell carcinoma. Oncogene 38:3475–3487.

Cannataro VL, Gaffney SG, Townsend JP. 2018. Effect Sizes of Somatic Mutations in Cancer. J Natl Cancer Inst 110:1171–1177.

Cannataro VL, Townsend JP. 2018. Neutral theory and the somatic evolution of cancer. Mol Biol Evol. doi:10.1093/molbev/msy079

Chapman AM, Sun KY, Ruestow P, Cowan DM, Madl AK. 2016. Lung cancer mutation profile of EGFR, ALK, and KRAS: Meta-analysis and comparison of never and ever smokers. Lung Cancer 102:122–134.

Doll R, Hill AB. 1950. Smoking and carcinoma of the lung; preliminary report. Br Med J 2:739–748.

Dubrow R, Darefsky AS. 2011. Demographic variation in incidence of adult glioma by subtype, United States, 1992-2007. BMC Cancer 11. doi:10.1186/1471-2407-11-325

Fousteri M, Mullenders LHF. 2008. Transcription-coupled nucleotide excision repair in mammalian cells: molecular mechanisms and biological effects. Cell Res 18:73–84.

Golemis EA, Scheet P, Beck TN, Scolnick EM, Hunter DJ, Hawk E, Hopkins N. 2018. Molecular mechanisms of the preventable causes of cancer in the United States. Genes Dev 32:868–902.

Greaves M. 2015. Evolutionary Determinants of Cancer. Cancer Discovery. doi:10.1158/2159-8290.cd-15-0439

Greenman C, Stephens P, Smith R, Dalgliesh GL, Hunter C, Bignell G, Davies H, Teague J, Butler A, Stevens C, Edkins S, O’Meara S, Vastrik I, Schmidt EE, Avis T, Barthorpe S, Bhamra G, Buck G, Choudhury B, Clements J, Cole J, Dicks E, Forbes S, Gray K, Halliday K, Harrison R, Hills K, Hinton J, Jenkinson A, Jones D, Menzies A, Mironenko T, Perry J, Raine K, Richardson D, Shepherd R, Small A, Tofts C, Varian J, Webb T, West S, Widaa S, Yates A, Cahill DP, Louis DN, Goldstraw P, Nicholson AG, Brasseur F, Looijenga L, Weber BL, Chiew Y-E, DeFazio A, Greaves MF, Green AR, Campbell P, Birney E, Easton DF, Chenevix-Trench G, Tan M-H, Khoo SK, Teh BT, Yuen ST, Leung SY, Wooster R, Futreal PA, Stratton MR. 2007. Patterns of somatic mutation in human cancer genomes. Nature 446:153–158.

Grolleman JE, Díaz-Gay M, Franch-Expósito S, Castellví-Bel S, de Voer RM. 2019. Somatic mutational signatures in polyposis and colorectal cancer. Mol Aspects Med 69:62–72.

Holford TR, Meza R, Warner KE, Meernik C, Jeon J, Moolgavkar SH, Levy DT. 2014. Tobacco control and the reduction in smoking-related premature deaths in the United States, 1964-2012. JAMA 311:164–171.

Hosseini S-R, Diaz-Uriarte R, Markowetz F, Beerenwinkel N. 2019. Estimating the predictability of cancer evolution. Bioinformatics 35:i389–i397.

Laconi E, Marongiu F, DeGregori J. 2020. Cancer as a disease of old age: changing mutational and microenvironmental landscapes. Br J Cancer 122:943–952.

Lawrence MS, Stojanov P, Polak P, Kryukov GV, Cibulskis K, Sivachenko A, Carter SL, Stewart C, Mermel CH, Roberts SA, Kiezun A, Hammerman PS, McKenna A, Drier Y, Zou L, Ramos AH, Pugh TJ, Stransky N, Helman E, Kim J, Sougnez C, Ambrogio L, Nickerson E, Shefler E, Cortés ML, Auclair D, Saksena G, Voet D, Noble M, DiCara D, Lin P, Lichtenstein L, Heiman DI, Fennell T, Imielinski M, Hernandez B, Hodis E, Baca S, Dulak AM, Lohr J, Landau D-A, Wu CJ, Melendez-Zajgla J, Hidalgo-Miranda A, Koren A, McCarroll SA, Mora J, Crompton B, Onofrio R, Parkin M, Winckler W, Ardlie K, Gabriel SB, Roberts CWM, Biegel JA, Stegmaier K, Bass AJ, Garraway LA, Meyerson M, Golub TR, Gordenin DA, Sunyaev S, Lander ES, Getz G. 2013. Mutational heterogeneity in cancer and the search for new cancer-associated genes. Nature 499:214–218.

Lee SG. 2016. Proving Causation With Epidemiological Evidence in Tobacco Lawsuits. J Prev Med Public Health 49:80–96.

Letouzé E, Shinde J, Renault V, Couchy G, Blanc J-F, Tubacher E, Bayard Q, Bacq D, Meyer V, Semhoun J, Bioulac-Sage P, Prévôt S, Azoulay D, Paradis V, Imbeaud S, Deleuze J-F, Zucman-Rossi J. 2017. Mutational signatures reveal the dynamic interplay of risk factors and cellular processes during liver tumorigenesis. Nat Commun 8:1315.

Liggett LA, DeGregori J. 2017. Changing mutational and adaptive landscapes and the genesis of cancer. Biochim Biophys Acta Rev Cancer 1867:84–94.

Li XC, Wang MY, Yang M, Dai HJ, Zhang BF, Wang W, Chu XL, Wang X, Zheng H, Niu RF, Zhang W, Chen KX. 2018. A mutational signature associated with alcohol consumption and prognostically significantly mutated driver genes in esophageal squamous cell carcinoma. Ann Oncol 29:938–944.

Loeb LA, Harris CC. 2008. Advances in chemical carcinogenesis: a historical review and prospective. Cancer Res 68:6863–6872.

Macintyre G, Goranova TE, De Silva D, Ennis D, Piskorz AM, Eldridge M, Sie D, Lewsley L-A, Hanif A, Wilson C, Dowson S, Glasspool RM, Lockley M, Brockbank E, Montes A, Walther A, Sundar S, Edmondson R, Hall GD, Clamp A, Gourley C, Hall M, Fotopoulou C, Gabra H, Paul J, Supernat A, Millan D, Hoyle A, Bryson G, Nourse C, Mincarelli L, Sanchez LN, Ylstra B, Jimenez-Linan M, Moore L, Hofmann O, Markowetz F, McNeish IA, Brenton JD. 2018. Copy number signatures and mutational processes in ovarian carcinoma. Nat Genet 50:1262–1270.

Martincorena I, Raine KM, Gerstung M, Dawson KJ, Haase K, Van Loo P, Davies H, Stratton MR, Campbell PJ. 2017. Universal Patterns of Selection in Cancer and Somatic Tissues. Cell 171:1029–1041.

Merlo LMF, Pepper JW, Reid BJ, Maley CC. 2006. Cancer as an evolutionary and ecological process. Nature Reviews Cancer. doi:10.1038/nrc2013

Montgomery ND, Selitsky SR, Patel NM, Neil Hayes D, Parker JS, Weck KE. 2018. Identification of Germline Variants in Tumor Genomic Sequencing Analysis. The Journal of Molecular Diagnostics. doi:10.1016/j.jmoldx.2017.09.008

Mroz EA, Tward AD, Hammon RJ, Ren Y, Rocco JW. 2015. Intra-tumor genetic heterogeneity and mortality in head and neck cancer: analysis of data from the Cancer Genome Atlas. PLoS Med 12:e1001786.

Nowell PC. 1976. The clonal evolution of tumor cell populations. Science 194:23–28.

Ostrow SL, Barshir R, DeGregori J, Yeger-Lotem E, Hershberg R. 2014. Cancer evolution is associated with pervasive positive selection on globally expressed genes. PLoS Genet 10:e1004239.

Petljak M, Alexandrov LB. 2016. Understanding mutagenesis through delineation of mutational signatures in human cancer. Carcinogenesis. doi:10.1093/carcin/bgw055

Pfeifer GP. 2015. How the environment shapes cancer genomes. Curr Opin Oncol 27:71–77.

Pfeifer GP, Denissenko MF, Olivier M, Tretyakova N, Hecht SS, Hainaut P. 2002. Tobacco smoke carcinogens, DNA damage and p53 mutations in smoking-associated cancers. Oncogene 21:7435–7451.

Poon SL, McPherson JR, Tan P, Teh BT, Rozen SG. 2014. Mutation signatures of carcinogen exposure: genome-wide detection and new opportunities for cancer prevention. Genome Med 6:24.

Poulos RC, Wong YT, Ryan R, Pang H, Wong JWH. 2018. Analysis of 7,815 cancer exomes reveals associations between mutational processes and somatic driver mutations. PLoS Genet 14:e1007779.

Ramakodi MP, Kulathinal RJ, Chung Y, Serebriiskii I, Liu JC, Ragin CC. 2016. Ancestral-derived effects on the mutational landscape of laryngeal cancer. Genomics 107:76–82.

Rawla P. 2019. Epidemiology of Prostate Cancer. World J Oncol 10:63–89.

Rosenthal R, McGranahan N, Herrero J, Taylor BS, Swanton C. 2016. DeconstructSigs: delineating mutational processes in single tumors distinguishes DNA repair deficiencies and patterns of carcinoma evolution. Genome Biol 17:31.

Segovia R, Tam AS, Stirling PC. 2015. Dissecting genetic and environmental mutation signatures with model organisms. Trends Genet 31:465–474.

Shield KD, Maxwell Parkin D, Whiteman DC, Rehm J, Viallon V, Micallef CM, Vineis P, Rushton L, Bray F, Soerjomataram I. 2016. Population Attributable and Preventable Fractions: Cancer Risk Factor Surveillance, and Cancer Policy Projection. Current Epidemiology Reports 3:201–211.

Shpak M, Lu J. 2016. An Evolutionary Genetic Perspective on Cancer Biology. Annu Rev Ecol Evol Syst 47:25–49.

Siegel RL, Miller KD, Jemal A. 2020. Cancer statistics, 2020. CA Cancer J Clin 70:7–30.

Smith MT, Guyton KZ, Gibbons CF, Fritz JM, Portier CJ, Rusyn I, DeMarini DM, Caldwell JC, Kavlock RJ, Lambert PF, Hecht SS, Bucher JR, Stewart BW, Baan RA, Cogliano VJ, Straif K. 2016. Key Characteristics of Carcinogens as a Basis for Organizing Data on Mechanisms of Carcinogenesis. Environ Health Perspect 124:713–721.

Somarelli JA, Gardner H, Cannataro VL, Gunady EF, Boddy AM, Johnson NA, Fisk JN, Gaffney SG, Chuang JH, Li S, Ciccarelli FD, Panchenko AR, Megquier K, Kumar S, Dornburg A, DeGregori J, Townsend JP. 2020. Molecular Biology and Evolution of Cancer: From Discovery to Action. Mol Biol Evol 37:320–326.

Stamatoyannopoulos JA, Adzhubei I, Thurman RE, Kryukov GV, Mirkin SM, Sunyaev SR. 2009. Human mutation rate associated with DNA replication timing. Nat Genet 41:393–395.

Sun W, Bunn P, Jin C, Little P, Zhabotynsky V, Perou CM, Hayes DN, Chen M, Lin D-Y. 2018. The association between copy number aberration, DNA methylation and gene expression in tumor samples. Nucleic Acids Res 46:3009–3018.

Temko D, Tomlinson IPM, Severini S, Schuster-Böckler B, Graham TA. 2018. The effects of mutational processes and selection on driver mutations across cancer types. Nat Commun 9:1857.

Tomasetti C, Li L, Vogelstein B. 2017. Stem cell divisions, somatic mutations, cancer etiology, and cancer prevention. Science 355:1330–1334.

Tomasetti C, Vogelstein B. 2015. Cancer etiology. Variation in cancer risk among tissues can be explained by the number of stem cell divisions. Science 347:78–81.

Townsend-Lab-Yale. n.d. Townsend-Lab-Yale/cancer_causes_and_effects. https://github.com/Townsend-Lab-Yale/cancer_causes_and_effects

Van den Eynden J, Larsson E. 2017. Mutational Signatures Are Critical for Proper Estimation of Purifying Selection Pressures in Cancer Somatic Mutation Data When Using the dN/dS Metric. Front Genet 8. doi:10.3389/fgene.2017.00074

Varley JM, Attwooll C, White G, McGown G, Thorncroft M, Kelsey AM, Greaves M, Boyle J, Birch JM. 2001. Characterization of germline TP53 splicing mutations and their genetic and functional analysis. Oncogene 20:2647–2654.

Venkatesan S, Birkbak NJ, Swanton C. 2017. Constraints in cancer evolution. Biochemical Society Transactions 45:1–13.

Wild C, Brennan P, Plummer M, Bray F, Straif K, Zavadil J. 2015. Cancer risk: role of chance overstated. Science.

Wu S, Powers S, Zhu W, Hannun YA. 2016. Substantial contribution of extrinsic risk factors to cancer development. Nature 529:43–47.

Yates LR, Campbell PJ. 2012. Evolution of the cancer genome. Nat Rev Genet 13:795–806.

Zhao Z-M, Zhao B, Bai Y, Iamarino A, Gaffney SG, Schlessinger J, Lifton RP, Rimm DL, Townsend JP. 2016. Early and multiple origins of metastatic lineages within primary tumors. Proc Natl Acad Sci U S A 113:2140–2145.

